# Wnt11 family dependent morphogenesis during frog gastrulation is marked by the cleavage furrow protein anillin

**DOI:** 10.1101/2022.01.07.475368

**Authors:** Elizabeth S. Van Itallie, Christine M. Field, Timothy J. Mitchison, Marc W. Kirschner

**Affiliations:** Department of Systems Biology, Harvard Medical School, Boston, MA 02115

**Keywords:** Gastrulation, Archenteron, Blastopore, Anillin, Non-canonical Wnt, Wnt11 family

## Abstract

Wnt11 family proteins are ligands that activate a type of Dishevelled-mediated, non-canonical Wnt signaling pathway. Loss of function causes defects in gastrulation and/or anterior-posterior axis extension in all vertebrates. Non-mammalian vertebrate genomes encode two Wnt11 family proteins whose distinct functions have been unclear. We knocked down zygotic Wnt11b and Wnt11, separately and together, in *Xenopus laevis*. Single morphants exhibited very similar phenotypes of delayed blastopore closure, but they had different phenotypes at the tailbud stage. In response to their very similar gastrulation phenotypes, we chose to characterize dual morphants. Using dark field illuminated time-lapse imaging and kymograph analysis, we identified a failure of dorsal blastopore lip maturation that correlated with slower blastopore closure and failure to internalize the endoderm at the dorsal blastopore lip. We connected these externally visible phenotypes to cellular events in the internal tissues – including the archenteron – by imaging intact embryos stained for anillin and microtubules. The cleavage furrow protein anillin provided an exceptional cytological marker for blastopore lip and archenteron morphogenesis and the consequent disruption through loss of Wnt 11 signaling. These cytological changes suggest a novel role for the regulation of contractility and stiffness of the epithelial cells that result in dramatic shape changes and are important in gastrulation.

## Introduction

Signaling mediated by Wnt11 family ligands is known to play an important role in gastrulation and anterior-posterior axis extension in all vertebrates that have been studied [1–7]. However, the specific requirement for these ligands during gastrulation has not been properly assessed by knock-down or knock-out perturbation in amphibians. Signaling downstream of Wnt11 family ligands is complex and involves multiple membrane receptors and cytosolic proteins [8, 9].

Except for the context of establishing the dorsal-ventral axis before gastrulation [10, 11], Wnt11 family ligands signal through non-canonical Wnt signaling pathways [6], which do not involve beta catenin. The best characterized non-canonical pathway is the Dishevelled-dependent “Planar Cell Polarity” (PCP) pathway. In this pathway Wnt11 family binding is thought to mediate the interaction of Dishevelled, the formin protein Daam1 [12], and RhoA resulting in increased Rho Kinase activity which mediates changes to the cytoskeleton [12–14]. JNK kinase activity is also regulated downstream of Dishevelled in this pathway [14]. Even though the endogenous requirements for the Wnt11 family ligands are not known, the results of disrupting PCP signaling in explants and in whole embryos have been well characterized using the dominant interfering Dishevelled construct Xdd1 [15–18]. This modification of Disheveled, where most of the PDZ domain is deleted, does not disrupt Adenomatous polyposis coli protein (APC) dependent degradation of beta-catenin in the canonical pathway [18]. The model amphibian *Xenopus laevis* played an important role in investigating the molecular biology of Dishevelled dependent PCP signaling, but the role of the Wnt11 family ligands specifically in morphogenesis of intact embryos has not been systematically investigated.

Non-mammalian vertebrate genomes typically encode two Wnt11 family proteins [19]. In *Xenopus laevis*, Wnt11b and Wnt11 have 63% identity at the protein level [20, 21] and are more similar to their presumed zebrafish orthologs than to each other (**Supplemental Figure 1**). In zebrafish, morpholino knockdown of either or both Wnt11 family proteins perturbs gastrulation, with evidence for partial redundancy [5]. In *Xenopus laevis*, a Wnt11b-based dominant negative construct initially established the importance of non-canonical Wnt11 family signaling during gastrulation and for anterior-posterior axis extension [6]. A translation blocking morpholino knockdown perturbation also identified a role for Wnt11b via the canonical Wnt signaling pathway in dorsal-ventral axis formation; additionally, a gastrulation defect phenotype was observed when the morpholino was injected after the dorsal-ventral axis was established [11, 22]. Wnt11b’s mRNA expression pattern in the pre-chordal mesoderm tissue during gastrulation is consistent with a gastrulation knock-down phenotype [6, 20]. Wnt11 has been studied in the context of Xenopus heart development [21, 23], but whether or not it has a role during gastrulation has not been investigated.

The three defining morphogenic events of gastrulation are the formation of the blastopore, the closure of the blastopore, and the formation of the archenteron. The blastopore is the circular structure on the surface of the vegetal pole of the embryo that will eventually be the anus. Surface ectoderm cells roll over the blastopore, forming an internal cavity, the archenteron; this proto anus-to-mouth cavity becomes the gut. Complex cell and tissue-level processes including bottle cell formation, convergent extension, and vegetal rotation are responsible for these structures. Bottle cells are epithelial cells that are extended in the apical-basal axis and have highly constricted apices. The pigment resulting from the concentration of the initially distributed pigment granules into the constricted apices is the first externally visible sign of gastrulation. Bottle cells are part of the early blastopore lip, and though they are not essential for blastopore closure, they are required for efficient closure [24, 25]. When the archenteron forms and extends, bottle cells are nearest the animal pole. Convergent extension is well-studied and is responsible for the ventral displacement of the blastopore as it closes. During convergent extension, cells intercalate along the medial-lateral plane, resulting in the extension of the tissue in the anterior-posterior axis [24,26,27]. Vegetal rotation is the dorsal and animal movement of the large endoderm cells near the vegetal pole of the embryo [28, 29]. Vegetal rotation generates animal movement of vegetal endoderm cells, which is important for the internalization of these cells as the blastopore closes. There are likely other cell-cell and tissue level processes that are important for the morphogenesis of these structures.

Here, we introduce anillin, a protein first known in cytokinesis and conserved from yeast to mammals [30], as a novel marker of morphogenesis in the blastopore lip and nascent archenteron. Anillin is best known as a cytokinesis furrow organizing protein which binds RhoA, F-actin, myosin II, and formins and membranes [30–35]. In its cytokinesis role, anillin binds active RhoA and stabilizes it at the cortex, and is necessary for stability of cleavage furrows and completion of cytokinesis [31]. It usually localizes exclusively to the nucleus in interphase cells, which is thought to provide a storage location prior to cytokinesis [36]. However, it was recently shown to localize to cell junctions in interphase cells of the Xenopus animal pole epithelium, where it promotes RhoA recruitment and contractility [37]. More recently, the effects of knockdown and overexpression of anillin were investigated at the tissue level of the animal pole epithelium. Perturbations to anillin levels were found to affect the amount of actomyosin that crosses the apical layer of cells resulting in changes to tissue stiffness and contractility [38]. The authors noted that anillin morphant embryos have a delayed gastrulation phenotype but did not explore this further [38]. Given the connection between anillin and regulation of cortical contractility by the RhoA pathway, as well as the reports of non-cytokinesis roles of anillin in epithelium, anillin was a candidate marker for epithelial shape change during gastrulation.

Using anillin as a marker, we report the phenotypes from knockdown perturbations of Wnt11b and Wnt11, individually and together, in whole embryos. We identify a phenotype of failed dorsal lip maturation, failed internalization of vegetal cells, and formation of the archenteron. We also show for the first time that anillin is an excellent marker for both bottle cells and the epithelial cells that undergo morphological changes in the early blastopore and later mature blastopore lip. These studies identify both cellular and tissue-scale features affected by perturbation of Wnt11 and focus on the transition from the surface epithelium to the formation of the archenteron.

## Results

### Wnt11b and Wnt11 function in blastopore closure

We analyzed recent transcriptomic [39] and proteomic [40] data to confirm the presence of Wnt11b and to look for evidence of Wnt11 expression during gastrulation in *Xenopus laevis* (**Figure 1**). Wnt11b mRNA and protein are detected during this period and increase during gastrulation. Wnt11 mRNA and protein levels are much lower than those of Wnt11b, but they also increase during this period. Since both Wnt11 family proteins are present during gastrulation, we investigated the phenotypes of knock-down perturbation of the proteins individually and together.

**Figure 1.**
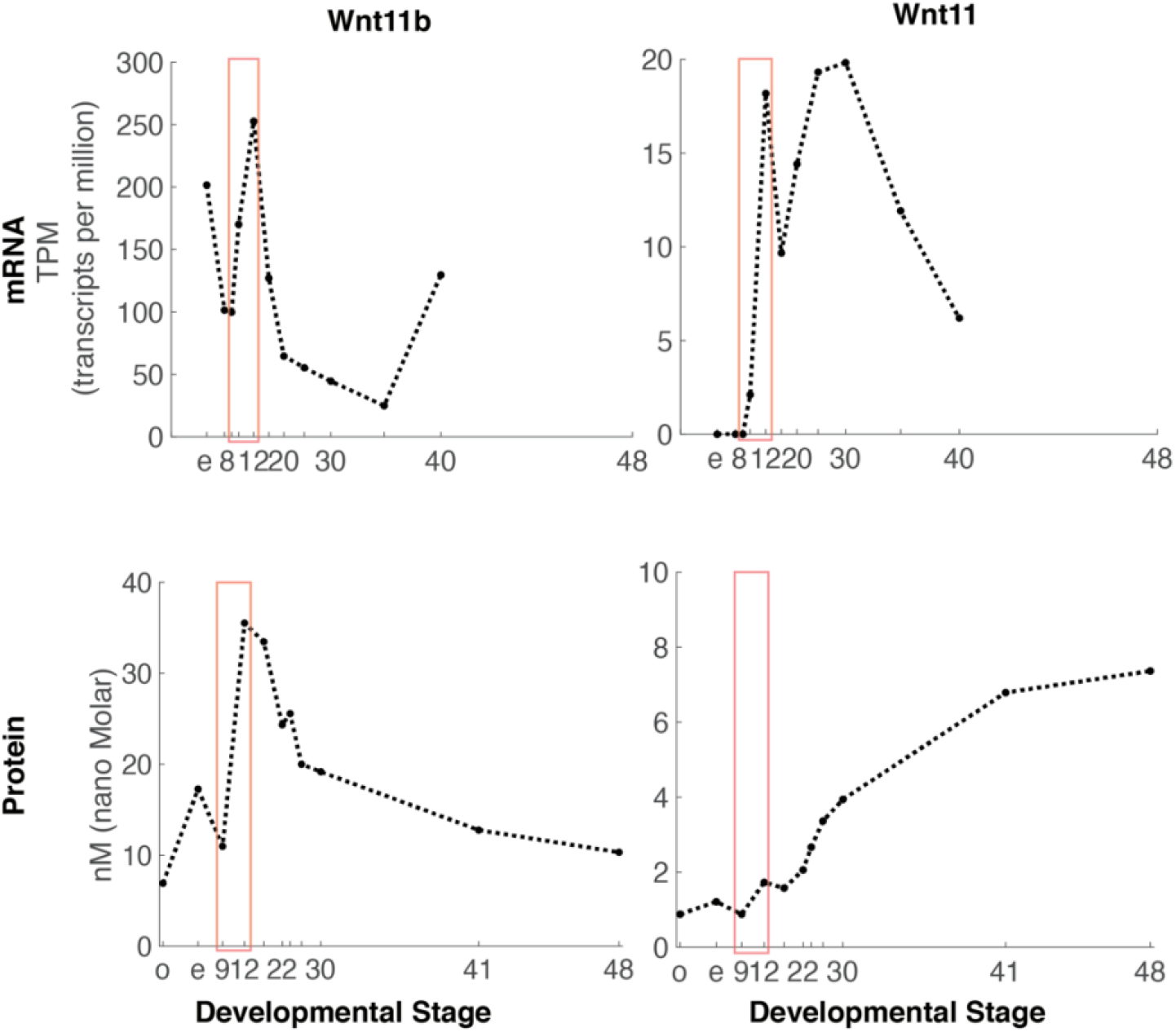
Recently published transcriptomic [39] and proteomic [40] data shows that both Wnt11b and Wnt11 increase in mRNA and protein level during gastrulation. Developmental time series transcriptomic and proteomic data both include multiple time points just before and during the gastrulation period (orange boxes; Stages 9,10, and 12 for transcriptomics and Stages 9 and 12 for proteomics). Note the much smaller y-axis scale for Wnt11 than Wnt11b for both mRNA and protein. There are two alloalleles of Wnt11: Wnt11.L and Wnt11.S. TPM counts for both Wnt11.L and Wnt11.S are combined. The peptides measured for Wnt11 do not allow us to distinguish between Wnt11.L and Wnt11.S.

**Figure 2.**
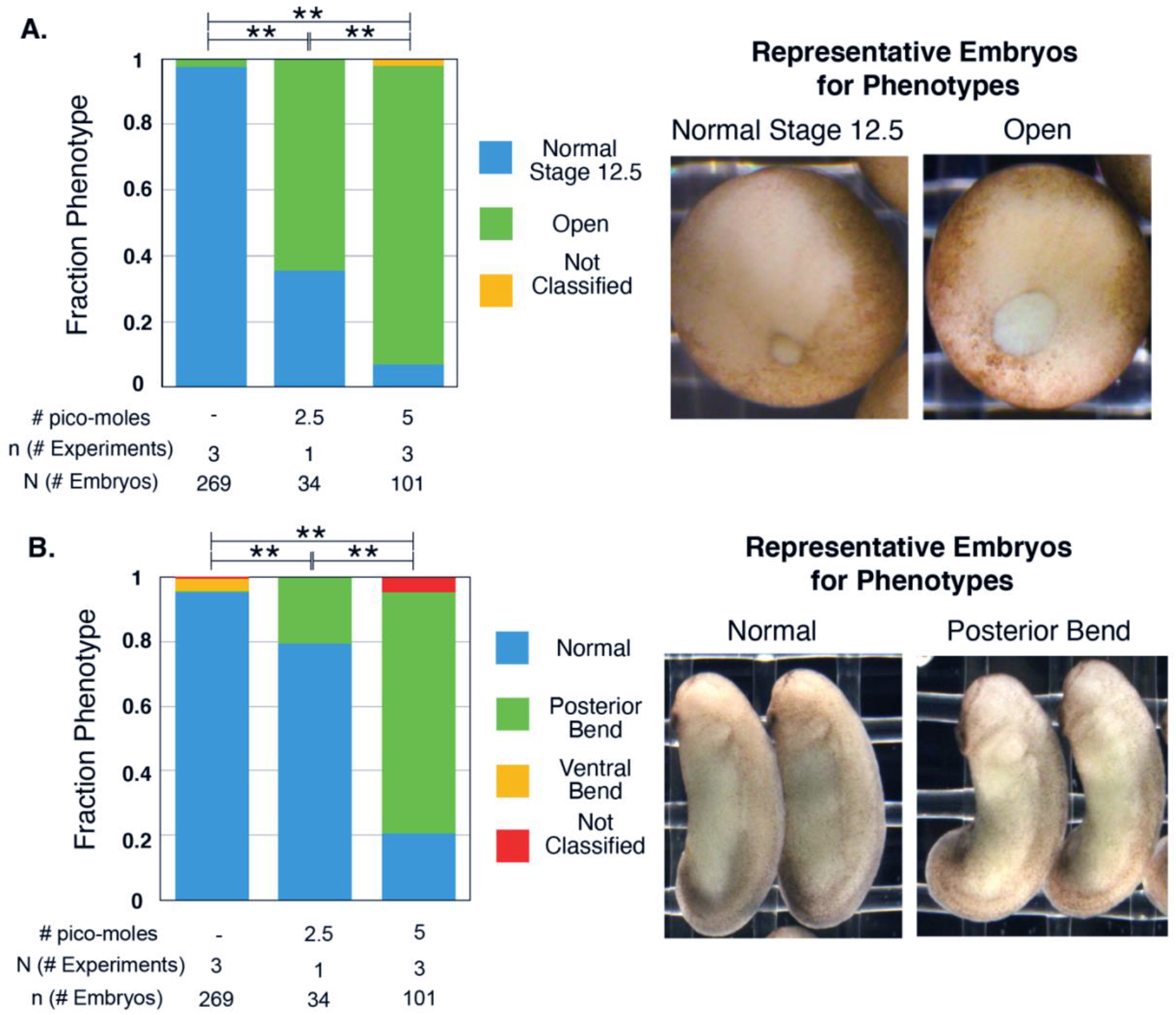
Wnt11b morphant embryos have larger blastopores at the end of gastrulation and posterior bends during the tailbud period. Embryos were injected with 2.5 or 5 picomoles of Wnt11b Mo [11, 22] total into both blastomeres at the 2-cell stage. The morpholino dose reported is the total dose. **A.** Injected and sibling control embryos were classified when sibling embryos are Stage 12.5 into two different phenotypes: normal and open blastopores. All conditions are statistically significantly different from each other (Fisher-Freeman-Halton test with BF method for multiple hypotheses; ** = p << 0.001). **B.** The same injected and sibling control embryos as in **A** were classified at the tailbud stage into three different phenotypes: normal, posterior bend, ventral bend (not shown). All conditions are statistically significantly different from each other (Fisher-Freeman-Halton test with BF method for multiple hypotheses; ** = p << 0.001).

**Figure 3.**
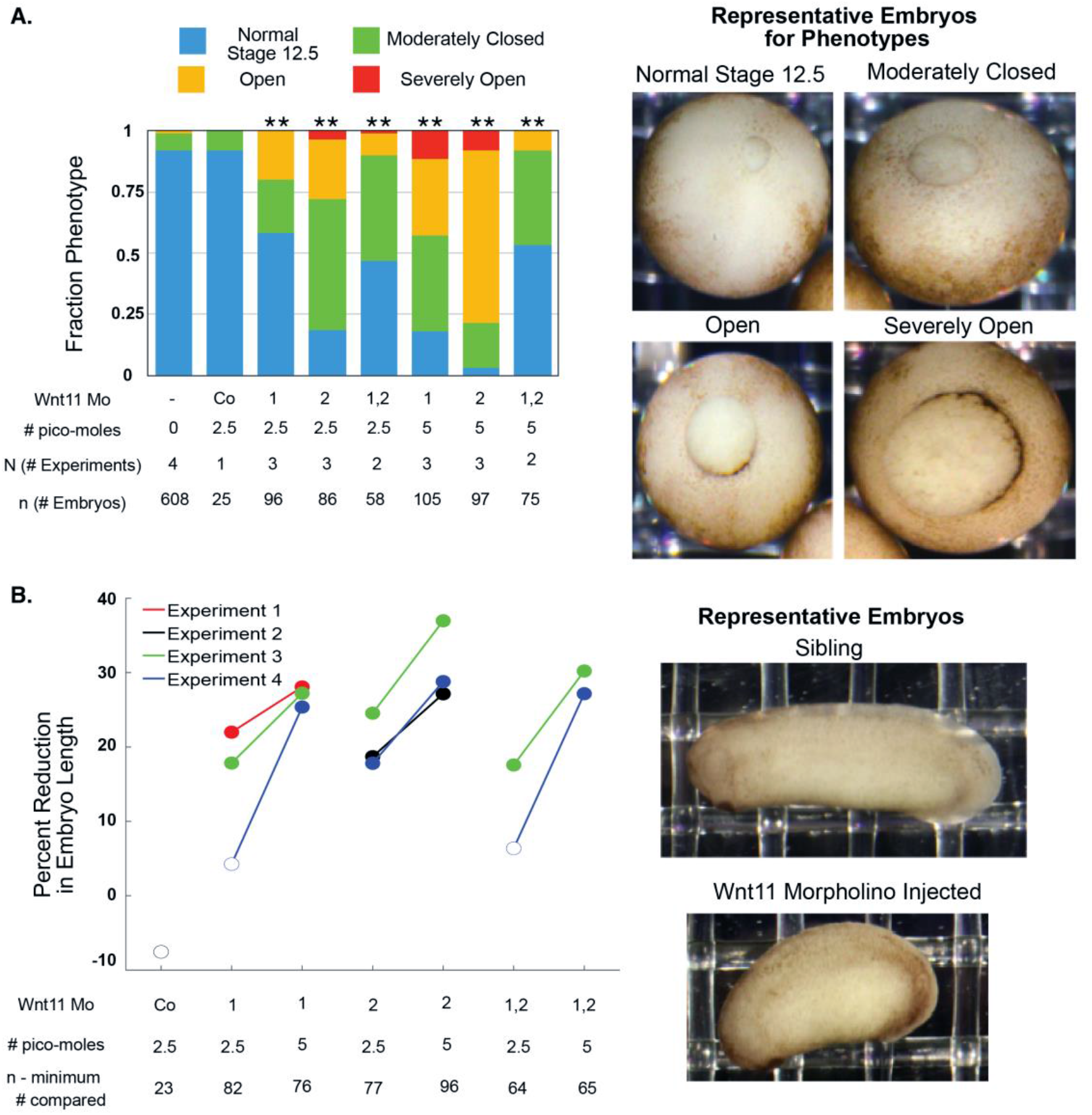
Wnt11 morphant embryos have larger blastopores at the end of gastrulation and shortened anterior-posterior axes during the tailbud period. Embryos were injected with 2.5 or 5 picomoles of Wnt11 Mo1, Wnt11 Mo2, or a 1:1 mixture of both into both blastomeres at the 2-cell stage. The morpholino dose reported is the total dose. **A.** Injected and sibling control embryos were classified when sibling embryos are at Stage 12.5 into four different phenotypes. All injections (except the standard control morpholino) resulted in statistically significantly different proportions of the phenotypes (Chi2 test; ** = p << 0.005). The phenotype for the 1:1 mixture of the two morpholinos is weaker than that of either of them alone. **B.** The anterior-posterior axis lengths of the same embryos as in **A** were quantified from still images at the tailbud stage. The statistical significance of the difference in the median length between the injected and sibling embryos was tested with the Wilcoxon Rank test using the Bonferroni method to correct for multiple hypothesis testing. The percent reduction of median length is plotted with the marker filled in if the difference is statistically significant (p < 0.05).

We first used a previously published Wnt11b translation-blocking morpholino to determine the resulting phenotypes from knockdown of Wnt11b protein levels [11, 22]. We injected both blastomeres at the 2-cell stage because Wnt11b is expressed circumblastoporally during gastrulation [6]. We found that Wnt11b morphant embryos exhibited more open blastopores at the end of gastrulation (**Figure 2.A**). The blastopore normally closes completely during neurulation (the morphogenic period following gastrulation where the neural folds appear and close and the anterior-posterior axis continues to extend). This also happened in the Wnt11b morphant embryos. There was no evidence of disruption of general features of the dorsal ventral asymmetry. However, when assayed much later during the tailbud stage, most Wnt11b morphant embryos had a posterior ventral bend (**Figure 2.B**). The anterior-posterior axis did not appear to be shorter than in un-injected siblings. Thus, we confirmed an apparently modest role of Wnt11b in gastrulation, based on its modest morphant phenotype.

We next considered the role of Wnt11. We confirmed the sequences of the 5’ UTRs of the transcripts of the two alloalleles for Wnt11 -- Wnt11.L and Wnt11.S -- and designed two different translation blocking morpholinos for each 5’ UTR. (**Supplemental Figure 2** for Wnt11.S morpholinos). Injection of either Wnt11.S morpholino at either of two different doses resulted in distinctly delayed blastopore closure (**Figure 3.A**). Since neither Wnt11.L morpholino gave a phenotype (not shown), we continued with only the Wnt11.S morpholinos and hereafter refer to Wnt11.S as Wnt11. As with Wnt11b morphants, the blastopores of almost all of the embryos eventually close during neurulation and there is no evidence of disrupted dorsal-ventral axis establishment (not shown). We also assessed the length of the anterior-posterior axis of the morphant embryos relative to un-injected siblings during the tailbud period (**Figure 3.B**.). In this case, the phenotype is different from the Wnt11b morphants. For all injections of the higher dose, morphant embryos have statistically significantly shorter anterior-posterior axes. There is no bend. The shortened anterior-posterior axis phenotype is consistent with in-situ hybridization evidence of Wnt11 expression in the somites during neurulation [21]. These data provide the first evidence for a role of Wnt11 that is similar to that of Wnt11b during gastrulation. After gastrulation, however, Wnt11’s role is different from that of Wnt11b.

When we injected a 1:1 combination of the two Wnt11 morpholinos, the blastopore closure phenotype was intermediate between the two single morpholinos at 2.5 picomoles and was not more severe at 5 picomoles (**Figure 3.A**). When the phenotype was assessed at the tailbud period, the effect from the 1:1 combination increased in severity with dose (**Figure 3.B**). Even though we did not observe a synergistic knockdown phenotype, we interpret the similarity in phenotype of the two morpholinos as evidence that the phenotype results from on-target, additive effects. These results establish a role for Wnt11 during gastrulation in *Xenopus laevis*.

Expecting redundancy in the roles of Wnt11 and Wnt11b during gastrulation, we perturbed both proteins simultaneously. We injected embryos with a 1:1 molar mixture of the Wnt11b morpholino and the Wnt11.S morpholinos (Wnt11b Mo: Wnt11 Mo1: Wnt11 Mo2 = 1:0.5:0.5). The total dose of morpholino injected was ten picomoles, compared to the maximum dose of five picomoles used to characterize the phenotype of the Wnt11b and Wnt11 morpholinos separately. As expected, Wnt11 family morphants have larger blastopores at the end of gastrulation compared to their matched sibling embryos, whereas control morphant embryos do not (**Figure 4**). These data are consistent with a partially redundant role of Wnt11b and Wnt11 during gastrulation. Therefore, we decided to pursue the double morphant approach to further characterize the role of Wnt 11 family signaling.

**Figure 4.**
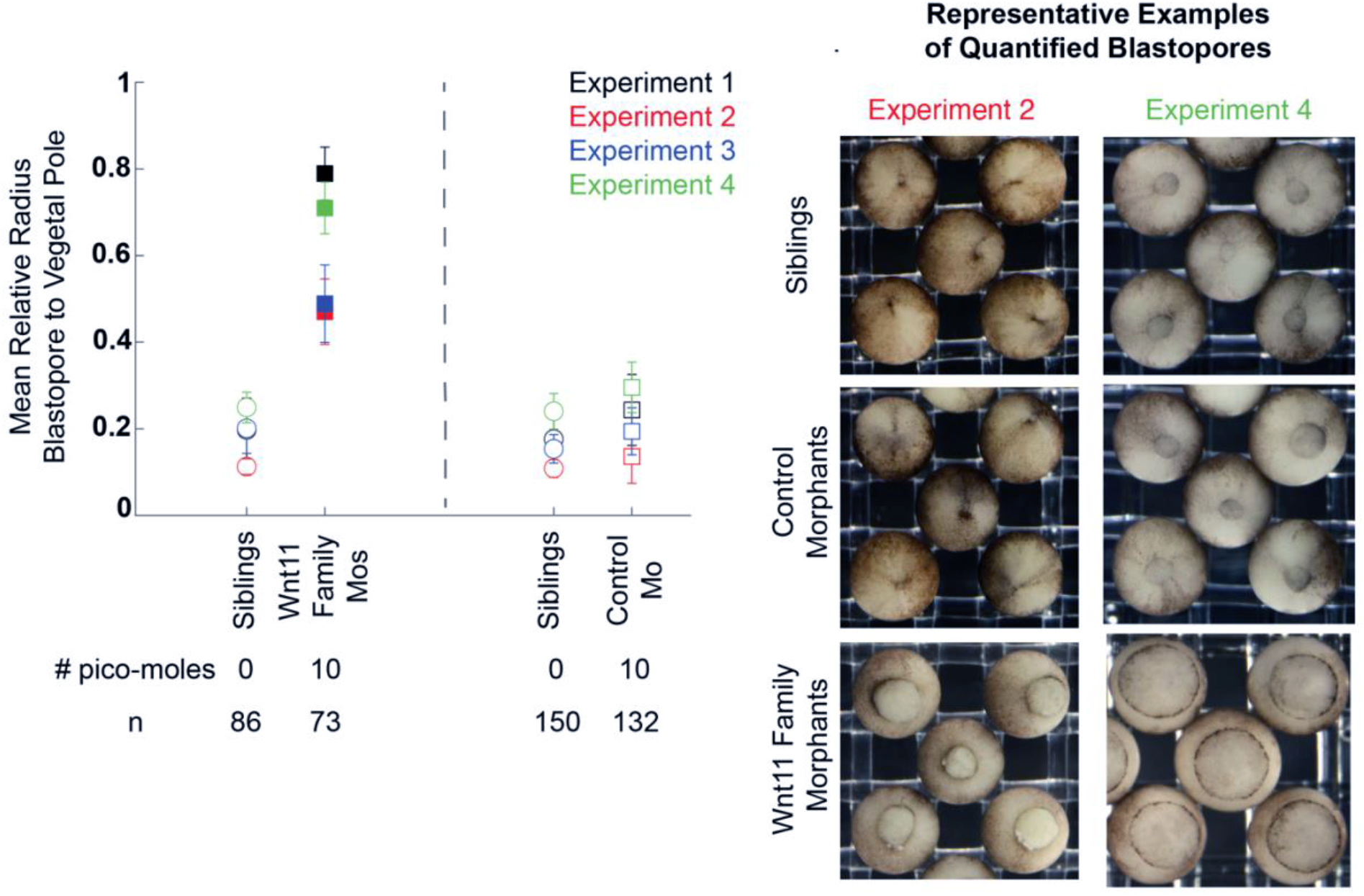
Wnt11 family morphants embryos have larger blastopores at the end of gastrulation than control morphants and siblings. For four biological replicates, embryos were injected with 10 picomoles total of either Wnt11 family morpholinos or the control morpholino in both blastomeres at the 2-cell stage. The relative radius of the blastopore to the embryo was quantified using FIJI processing of still images taken when siblings embryos were at Stage 12.5. The difference in relative blastopore radius was statistically significant for all replicates. The sample mean and standard deviation are shown for all biological replicates and experimental conditions. Statistical significance of the difference between the experimental injections and standard control morpholino injection values for each clutch was determined with the student’s t-test, and statistically significant perturbation have filled in markers. Multiple hypothesis testing was corrected for using the Bonferroni Method. Representative example blastopores from the different conditions are shown for two biological replicates.

### Time lapse imaging of blastopore closure

To characterize the Wnt11 family morphant phenotypes with respect to the external events of gastrulation, we performed multiplexed time-lapse imaging of the vegetal pole during gastrulation. Dark-field illumination revealed cell boundaries of the blastopore lip with excellent contrast. Up to 24 embryos were imaged per experiment, allowing sibling (un-injected) embryos to be used to control for imaging conditions and timing. A typical un-injected embryo is shown in **Movie 1**, a control morphant in **Movie 2**, and Wnt11 family morphants in **Movie 3** and **Movie 4**. Inspection of the movies showed that the Wnt11 family morphant blastopores closed more slowly, as expected from the gastrulation endpoint results, and also that the dorsal blastopore lip is not capable of internalizing the dorsally moving vegetal cells efficiently. As a result, the dorsally moving vegetal material “overflows” the dorsal blastopore lip. Furthermore, the blastopore lip often appeared mechanically unstable in Wnt 11 family morphants, with evidence of the ring diameter transiently increasing.

To compare and visualize dynamic features of blastopore formation and closure across the dataset, we made kymographs from the image stacks where the distance axis was a line bisecting the embryo passing first through the dorsal lip and then across the later-forming ventral lip (**Figure 5**). These kymographs capture the appearance and movement of the dorsal and ventral blastopore lips across the embryo. The closer the kymograph line is to horizontal, the faster the movement. We classified the kymographs into three phenotypes: “normal,” “mildly perturbed,” and “severely perturbed.” In the “normal” kymographs, as time progresses, the dorsal blastopore lip line appears with a dark pigmentation and moves towards the center of the kymograph at roughly constant velocity. Later, the ventral lip pigmentation appears and also moves towards the center of the kymograph (**Figure 5.A.i**). Then, the initially dark dorsal lip line becomes lighter in color when the bottle cells, associated with much of the pigment, have moved inside the embryo. The dorsal lip line continues to smoothly move towards the ventral side of the kymographs, and the ventral line continues to move dorsally. This decreasing distance between the dorsal and ventral lip lines is the kymograph representation of the blastopore closing.

**Figure 5.**
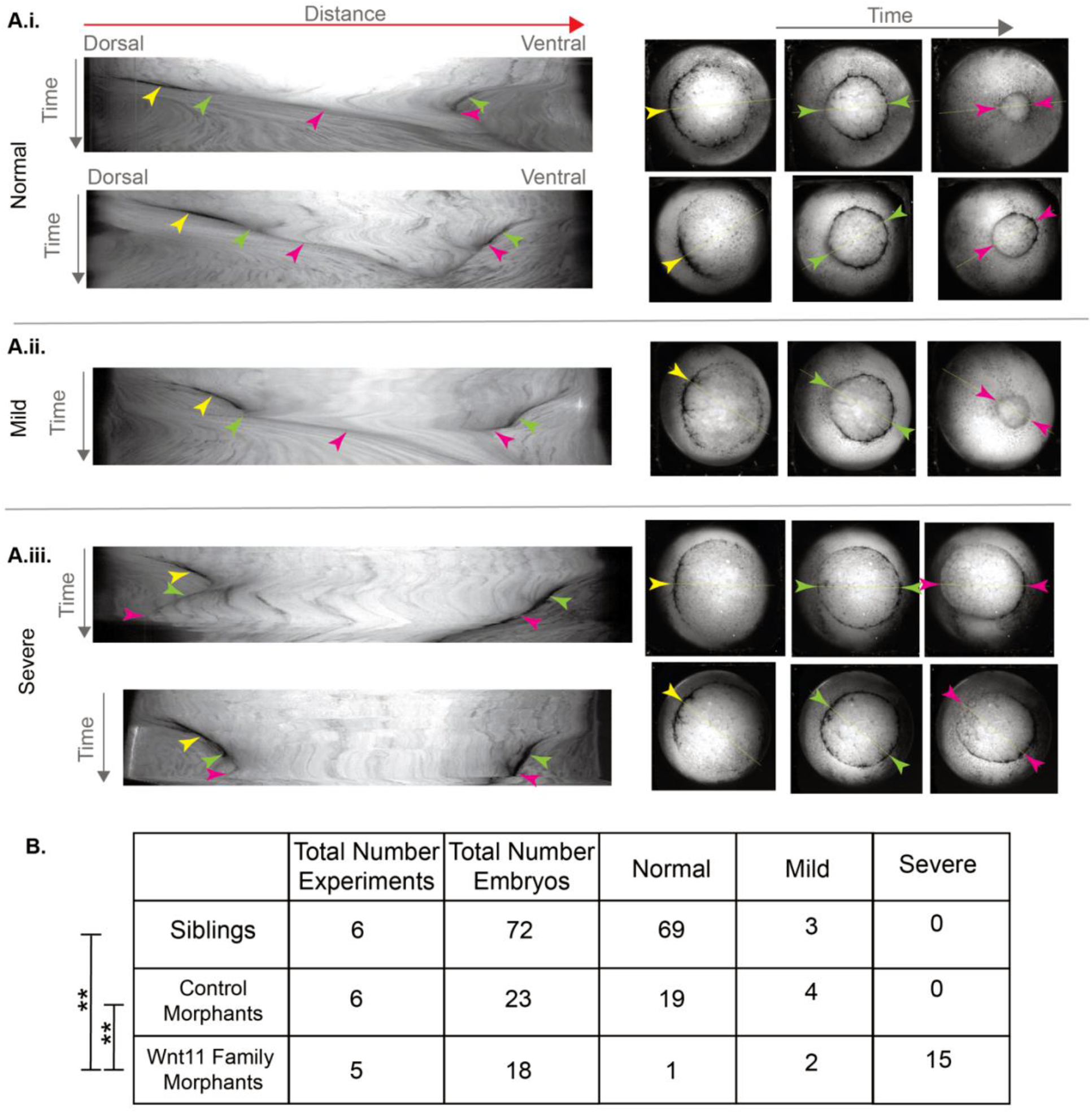
Time lapse imaging reveals blastopore maturation, bottle cell internalization, and blastopore closure phenotypes associated with the Wnt11 family morphants. **A.** Kymographs were made from the time lapse images of the vegetal pole during gastrulation. The distance axis is the line across the vegetal pole that passed first through the center of the dorsal lip and then through the center of the eventual ventral blastopore lip. The kymographs were classified into “normal,” “mild perturbation,” and “severe perturbation.” Representative examples of the three classes are shown. In the kymographs for all three phenotypes the dorsal bottle cells appear and move towards the center of the embryo and an early lip has formed (yellow arrowhead). In the “severe perturbation” kymographs the movement towards the center is slower (compare **i.** and **ii.** to **iii.**). In the normal kymographs (**i.**), after the blastopore has moved further, the early lip matures and the color of the dorsal lip line changes to become lighter because the bottle cell have moved inside the embryo (dorsal green arrow). In the mild kymographs (**ii.**), this happens after transient reversal in the movement of the dorsal lip (dorsal green arrow). In the kymographs for all three phenotypes, the dark ventral lip line appears and also moves towards the center of the embryos (ventral green arrow). In the normal and mild kymographs, the dorsal and ventral lip continue to move closer to each other (pink arrowheads). By contrast, in the severe kymographs, the dorsal lip either reverses and moves away from the ventral line or stalls (dorsal pink arrowhead). **B.** Results from qualitatively scoring the kymographs from a total of five (Wnt11 family morphants) and six (sibling and control morphants) independent experiments. The Wnt11 family morphants are statistically significantly different than the control morphants and siblings (Fisher-Freeman-Halton test with BF method for multiple hypotheses; ** = p << 0.001).

In the “severely perturbed” kymographs, which constituted 15/18 (83%) of the Wnt11 family morphant data, the dorsal and ventral lip pigmentation lines also appear and initially move towards the center of the kymograph. However, the movement of the dorsal blastopore lip is slower than in the normal kymographs, and the dark pigment color is not lost because the bottle cells with their associated pigment do not involute. Eventually, the dorsal lip line either stops moving or starts moving dorsally, i.e. the reverse of its normal ventral movement (**Figure 5.A.iii**). When the movies corresponding to these kymographs are inspected, we see that the dorsally moving vegetal cells have overflowed the dorsal blastopore lip instead of being internalized (**Movie 3** and **Movie 4**). Thus, the “severely perturbed” kymographs reveal the disruption of an important transition event during gastrulation: the change of the dorsal lip from one with bottle cells on the surface to one where they have been internalized. We consider the dorsal lip after the bottle cells have been internalized to be a “mature lip.” Without a mature lip, the embryos do not appear capable of internalizing the dorsally moving vegetal material.

We scored 2/18 (11%) of the Wnt family morphant and 4/23 (17%) of control morphant kymographs as “mildly perturbed” (**Figure 5.B**). In these kymographs a small stall or reversal of the dorsal lip line occurred but the dorsal lip eventually matured and moved ventrally (**Figure 5.A.ii**). Based on the classification of kymographs into the three different categories, the Wnt11 family morphants are statistically significantly different than the siblings and control morphants.

Eighty-three percent of the Wnt11 family morphants have the severe phenotype compared to none of the sibling or control morphants (**Figure 5.B**).

We also quantified the relative radius of the blastopore compared to that of the embryo for all of the time lapse movies starting at the time point when the matched sibling embryos have just formed mature lips and internalized the bottle cells (**Supplemental Figure 3**). As expected from the qualitative inspection of the movies, the Wnt11 family morphants have larger blastopore radii at this timepoint than the sibling embryos and control morphants. Then, during the time period that the siblings and control morphants completely close their blastopores, Wnt11 family morphants show only minimal blastopore closure. Taken together with the kymograph analysis, Wnt11 family morphants have decreased initial movement of the dorsal blastopore lip, failure to form a “mature” blastopore lip, failure to internalize the dorsally moving vegetal material, and extremely reduced blastopore closure. However, knock-down of Wnt11 family proteins does not perturb bottle cell formation or dorsal flow of the vegetal cells.

### Cortical anillin marks epithelial cells in the early blastopore lip

Next, we used immunofluorescence to assess the internal structures of the embryo during the periods before and after the Wnt11 family dependent failure of blastopore lip maturation. We sought marker proteins for visualizing the cytoskeleton during these periods. Our fixation and clearing methods made visualization of F-actin difficult. We found that microtubules were a good marker of cell boundaries and that the cleavage furrow protein anillin provided exceptional record of epithelial cells undergoing shape change. Both tubulin and anillin have well-established localizations during mitosis and cytokinesis, which we confirmed in dividing cells during gastrulation (**Supplemental Figure 4).** These observations helped validate our fixation methods and antibodies.

Anillin exhibited dramatic enrichment at the contracting apex of bottle cells, as they initiate blastopore lip morphogenesis at Stage 10.5. **Figure 6.A** shows an animal-vegetal cross-section view; see **Supplemental Figure 5.A** for a schematic relating this view to the exterior vegetal view of the whole embryo. Anillin was also enriched on the outside cortex of pre-bottle cells before lip invagination. **Figure 6.B** shows multiple dorsal-ventral cross-section views of the forming dorsal blastopore lip. In the most vegetal section, anillin is punctate and scattered along outer membranes of the cells that are changing shape. In more animal planes, anillin forms a contiguous line along the outer membranes of the cells. Since the forming blastopore lip is moving vegetally, we assume that the contiguous line is the temporally later localization. We do not know the mechanistic relevance of these two different localizations. In the dorsal-vegetal cross-section view we can see that not all of the cells at the forming lip have constricted apices in this plane. Some cells are elongated, and they have the membrane proximal anillin staining. It is known that elongated cells that are not apically constricted are also part of the blastopore lip [41, 42]. Anillin appears to be in a membrane proximal location in cells that are changing shape.

### Wnt11 family morphants make an unsuccessful attempt to form a dorsal lip

With anillin as a marker for the progression of lip morphogenesis, we were able to investigate the defects in Wnt11 family morphants in more detail. First, we wanted to determine how the early lip compares between siblings and Wnt11 morphants during the period before we see the phenotype of “failed lip maturation.” We imaged sibling, control morphant, and Wnt11 morphants fixed at the same timepoint and classified the resulting immunofluorescence images into “normal lips,” “abnormal lips,” or “no lip” (**Figure 7, top lef**t). All of the sibling and control morphant embryos form normal early lips (**Figure 7, top right**). By contrast, more than 80% of Wnt11 family morphants either have an abnormal lip or no lip at all (**Figure 7, bottom**). In all phenotypes there are cells that have membrane proximal anillin and this anillin localization is only on the dorsal side of the embryo. This is consistent with the fact that all Wnt11 family morphants form bottle cells and that dorsal-ventral axis formation is not perturbed by the morpholinos. In embryos with “abnormal lips,” we see arcs of elongated cells that have membrane proximal anillin, but the lines of anillin are more spotted and are thinner. The abnormal lip structure is also closer to the edge of the embryo. The “no lip” phenotype has puncta but no lines of anillin and also is close to the edge of the embryo.

**Figure 7.**
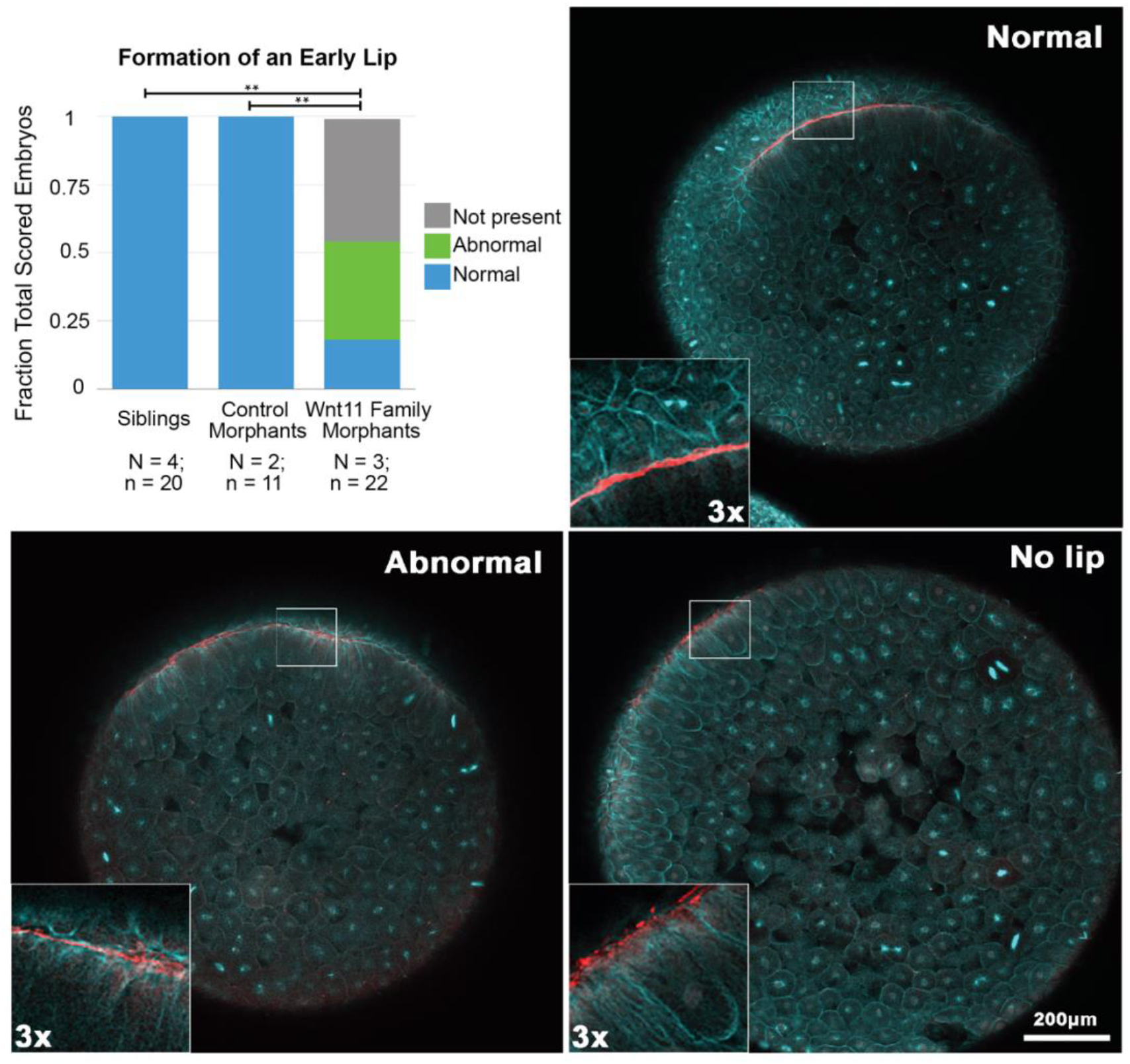
Formation of an early blastopore lip is perturbed in Wnt11 family morphants. Stacked bar charts showing the results of scoring formation of an early lip in sibling, control morphant, and Wnt11 family morphant embryos across multiple independent experiments (top left). For both scored phenotypes, the Wnt11 family morphants are statistically significantly different from the sibling and control morphant embryos (** = p<< 0.05). Immunofluorescence panels are representative examples of the normal, abnormal, and not present phenotypes scored.

Our immunofluorescence data of unperturbed embryos revealed that as the lip forms the anillin staining changes from spotted (**Figure 6.B, left**) to solid lines (**Figure 6.B, center and right**). The “no lip” phenotype shows the spotted localization, but there is no evidence of further progression. The “abnormal lip” phenotype does have lines of anillin, but they are not as continuous or as thick as those of normal lips. The fact that greater than 80% of Wnt11 family morphants have “abnormal” or “not present” lips is evidence that Wnt11 family morphant embryos can take the first early steps to make lips, but these early lips are usually not reinforced and thus the lips fail to develop properly.

### The formation of a nascent archenteron after blastopore lip maturation

Anillin and tubulin provided excellent markers to investigate the properties of the dorsal blastopore lip after lip maturation (Stage 11). We fixed sibling embryos during this period to determine the normal internal phenotype. **Figure 8.A** shows an animal-vegetal cross-section view of anillin and tubulin staining (see **Supplemental Figure 5.B** for schematic). The dorsal bottle cells have moved inside the embryo and the former suprablastoporal (above the bottle cells) and subblastoporal (below the bottle cells) endoderm now line the archenteron. These cells that line the archenteron have anillin proximal to the plasma membrane (**Figure 8.A, left inset**). On the ventral side of the embryo the bottle cells and ventral blastopore lip are caught in the process of formation (**Figure 8.A, right inset**).

**Figure 8.B** shows three z-planes of the dorsal-ventral view of mature lip and upwardly moving archenteron from an embryo. The shapes of the cells that have anillin staining and of the anillin staining itself are different in the three panels. In the more vegetal plane, we see that the cells of the blastopore lip are columnar and show organized packing. In this optical plane, anillin has a smooth membrane proximal localization (inset below). As we move upward, towards the animal pole, the shape of the cells and the anillin staining change. The cells become less and less columnar, and the anillin staining becomes rougher (center). We also see that there are two lines of anillin staining with a small gap between them. This gap can also be seen in the tubulin staining, which clearly reveals cell shape. We appear to be viewing both sides of the epithelium that lines the slit-like archenteron. In the plane closest to the animal pole, the cells are the most spherical in shape and the gap between the two sides is smaller (right). When we compare the whole embryo views in the different planes, we see that the span of the anillin staining shortens as the planes move animally. This shows that during this period of gastrulation the archenteron narrows in the animal pole direction and bottle cells are at the animal extent.

### Wnt11 family morphants exhibit defective archenteron extension

To investigate the role of Wnt11 family signaling in morphogenesis of the mature blastopore lip and nascent archenteron, we fixed siblings, control morphants, and Wnt11 family morphants and scored archenteron extension using anillin and tubulin staining (**Figure 9**). All of the sibling and control morphant embryos have normal archenteron extension (**Figure 9.A**; see **Figure 9.B** for representative example). However, ∼90% of Wnt11 family morphants show no evidence of archenteron extension. We show two representative examples of embryos with no archenteron formation. In these embryos we do not see the overlaying arcs of anillin as the planes more animally or any evidence of a short arc of bottle cells at the animal extent of the planes of membrane proximal anillin staining. In one of the no-archenteron examples (**Figure 9.C.i**), we do see a successfully formed immature lip (orange asterisks). This is the only example in the data where we see a successful immature lip but no archenteron formation. **Figure 9.C.ii** has multiple foci of membrane proximal anillin (yellow asterisks), but they have not progressed to a correctly formed immature lip. Interestingly, the arc that spans these foci is wider than what was observed in the earlier images Wnt11 morphants attempting to form a lip. This observation is consistent with Wnt11 family morphant also forming laterally and eventually ventral bottle cells. In the normal archenteron formation example, we see larger foci of anillin and bottle cells opposite the forming archenteron which is evidence of the normally forming ventral lip.

**Figure 9.**
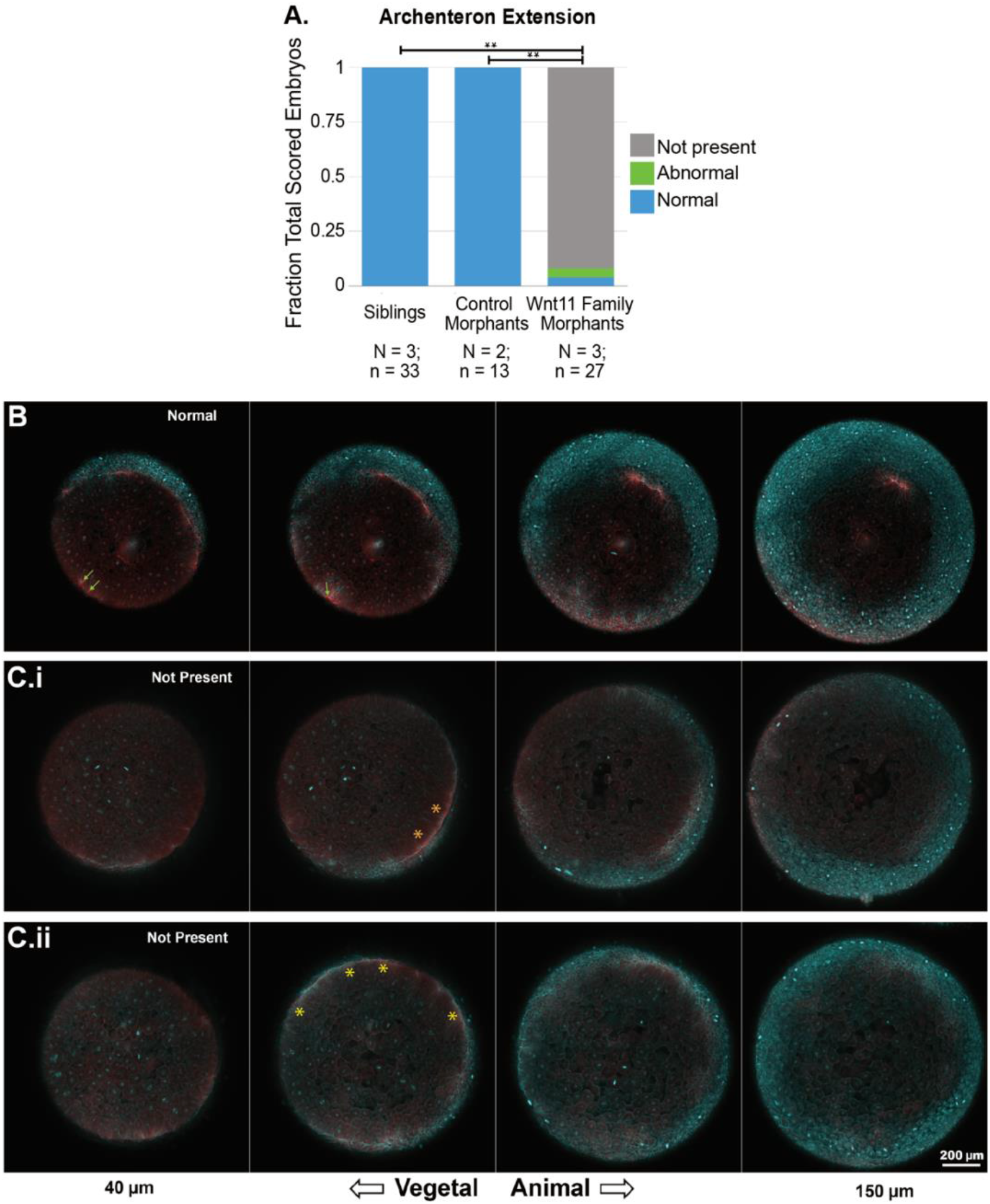
Wnt11 family morphants exhibit no archenteron extension. **A.** Stacked bar charts showing the results of scoring archenteron extension in sibling, control morphant, and Wnt11 family morphant embryos across multiple independent experiments. In siblings and controls, the archenteron is present in all embryos. By contrast, in greater than 90% of Wnt11 family morphants, the archenteron is **not** present. Wnt11 family morphants are statistically significantly different from the sibling and control morphant embryos (** = p<< 0.05). **B.** A representative example of the archenteron staining seen in sibling and control morphants after lip maturation. Anillin is present in multiple dorsal-ventral planes spanning over 100 μM. The span of the anillin staining gets shorter as the planes move through the embryo animally. Green arrows point to foci of anillin and bottle cells opposite the archenteron staining that signal the forming ventral lip. **C.** Representative examples of the observed failure of archenteron formation seen in the vast majority of Wnt11 family morphants. In both examples, we see membrane proximal anillin staining in one of the planes (asterisks) but neither of the more animal planes; i.e. there are no overlaying and shortening arcs of anillin across multiple planes. **C.i.** is one of the rare examples where there is evidence of a successfully formed immature lip. **C.ii.** is much more representative of the “not present” phenotype. There are multiple membrane proximal anillin foci, but they have not formed a lip with lines of anillin.

## Discussion

Amphibian embryos have always played a large role in understanding the morphogenesis that takes place between the spherical egg, a geometry shared by many organisms, and the vertebrate body plan [43]. The initial starting conditions of egg shape and yolk distribution, which vary greatly across vertebrates, are important to the strategies of gastrulation [44], but there is evidence that some of the cell and tissue level behaviors, as well as the signaling pathways that control them, are conserved [45, 46]. One of these widely used signaling pathways is Dishevelled dependent non-canonical Wnt signaling. We have learned a great deal about the molecular players involved in non-canonical Wnt signaling, but we still have an imperfect idea of how a process like gastrulation use these pathways at the tissue level.

Our study began with an attempt to understand the roles of the two Wnt11 family proteins Wnt11 and Wnt11b, thought to be agonists in the non-canonical Wnt pathway. In what we report here, this investigation leads to very fundamental questions, such as how the blastopore forms, how the vegetal endoderm is internalized, and how the archenteron forms and extends. We introduced two new methods: multiplexed darkfield movies of vegetal pole dynamics, and whole mount immunofluorescence with antibodies to tubulin and anillin. The movies of the vegetal pole led to the finding that Wnt11 family proteins signal to promote blastopore lip maturation, internalization of the vegetal material, and blastopore closure. Whole mount immunofluorescence revealed strong enrichment of anillin in the outer cortex of epithelial cells in the blastopore lip and archenteron. We were able to use anillin as a cytological marker for epithelial dynamics in the gastrula, revealing that Wnt11 family signaling is also involved in archenteron formation. All of these phenotypes (blastopore lip formation, vegetal material internalization, blastopore closure, and archenteron formation) have been reported in *Xenopus laevis* embryos perturbed with dominant negative Dishevelled, which provides strong evidence that the Wnt11 family signaling during gastrulation is Dishevelled dependent [15]. Although some of these events are due to the behavior of the mesoderm and convergent extension, the behavior of epithelial cells is also affected. Our findings suggest, along with those of others [47], suggests that these epithelial cells are generating their own forces, as well as stabilizing forces generated by the tissues around them. We found that not only was the tissue organization of the dorsal blastopore lip very different in the sibling and Wnt11 family morphants, but the anillin staining was also very different. This of course suggests that anillin’s role in maintaining tension and structural integrity of the lip is dependent on Wnt11 family signaling. But it also suggests that imaging of other cellular components in control and Wnt11 morphants will further increase our mechanistic understanding. Studies in whole embryos are particularly important in gauging the activities coordinated in space and time that will be necessary to understand the effects of signaling on the mechanics of gastrulation as a whole; it supplements information previously gained from explant cultures [6,18,27].

This is the first experimental paper, to our knowledge, that has explicitly addressed the function of the two Wnt11 family members in *Xenopus laevis*. We point out that there are inconsistencies in the names used to refer to Wnt11b and Wnt11 in the Xenopus literature, which has added confusion. Our results show that Wnt11, in addition to Wnt11b, is involved in gastrulation in *Xenopus laevi*s. We have not directly addressed the overlapping versus non-overlapping roles of Wnt11b and Wnt11 during gastrulation or later periods in embryogenesis. In-situ hybridization data of Wnt11b and Wnt11, alone and together, during gastrulation in whole and bisected embryos would add to our knowledge of the roles of these proteins. The mRNA and protein levels of Wnt11b and Wnt11 are notably very different, with Wnt11 being much lower. It would be interesting to quantify the effects of the knockdowns on the levels of Wnt11b and Wnt11 to assess how absolute protein amounts affect the phenotypes. In addition, there is a third non-canonical Wnt ligand in the Xenopus laevis genome: Wnt5a. The phenotype of knockdown perturbation of Wnt5a in whole embryos has never been reported. Tada & Smith reported that Wnt5a and Wnt11b rescued dominant negative Wnt11b disruption of Dishevelled phosphorylation [6]. Based on this finding, if both proteins were expressed in the same tissues, redundancy would be expected. Our Wnt11 family knockdown perturbation would be useful for determining the extent to which Wnt5a signaling contributes to morphogenesis in *Xenopus laevis*.

The work reported here provides new information on an under-studied problem: morphogenesis of the early archenteron in vertebrates, a critical component of the vertebrate body plan [48, 49]. Cortical anillin is strongly enriched in the epithelial cells that will line the archenteron (**Figures 6**, **8**). How the initial bottle cells mature into a lip and then reorganize in the animal-vegetal direction to form an extended groove is not known. One hypothesis for their extension animally after involution is that the bottle cells are pulled toward the animal pole by either the involuted mesoderm or migrating head mesoderm [25, 49]. Ewald et al. found that perturbation of Dishevelled signaling also results in decreased archenteron extension [15]. With our results, there is now strong evidence that archenteron extension requires Wnt11 family non-canonical Wnt signaling. But are these epithelial cells responding to Wnt11 family ligands themselves?

**Figure 6.**
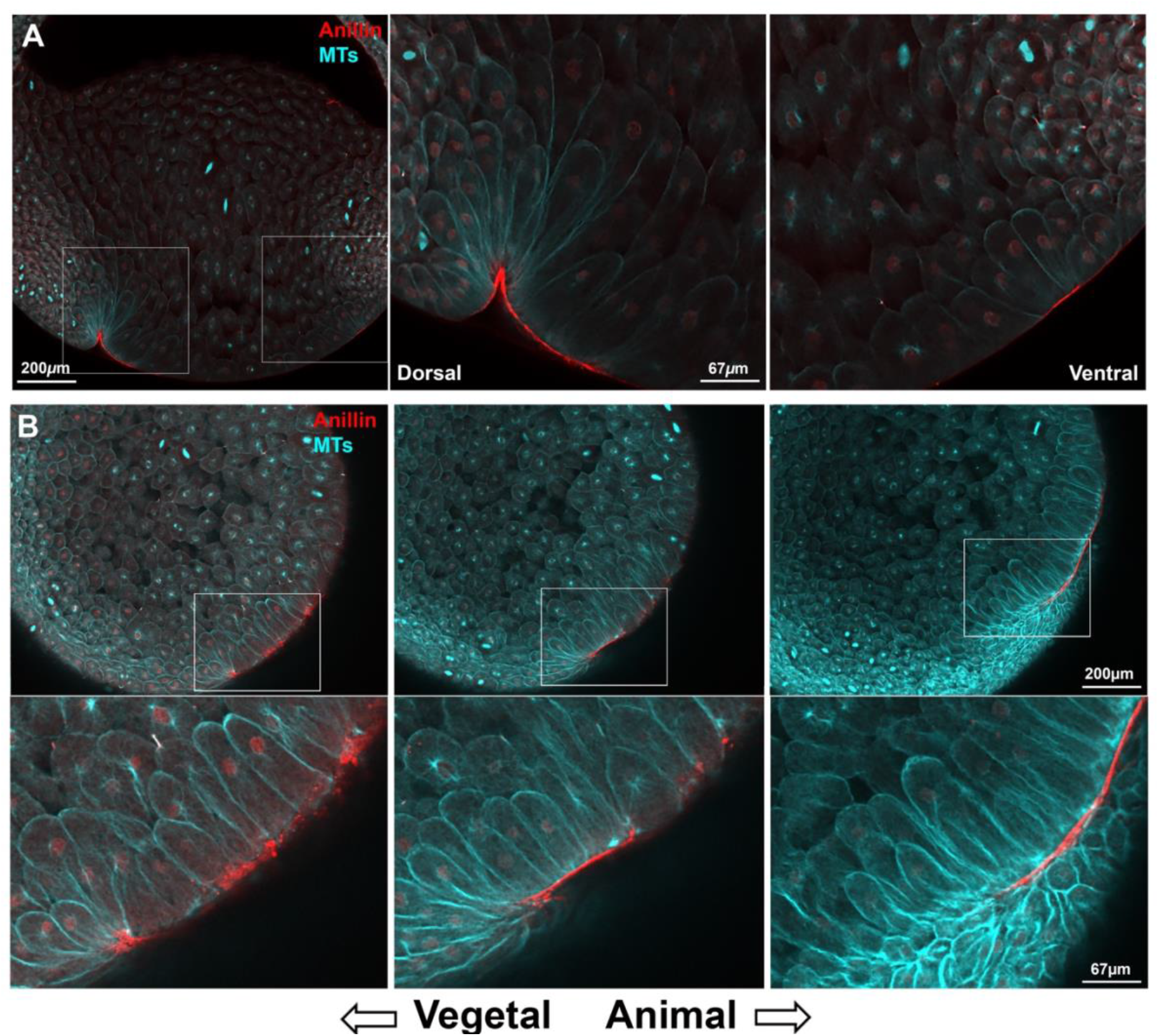
Whole embryo fixed immunofluorescence against tubulin and anillin during the formation of the dorsal lip. **A.** Animal-vegetal optical plane view of a Stage 10.5 embryo shows membrane proximal anillin localization to bottle cells and pre-bottle cells. Tubulin staining shows the cell outlines, revealing bottle cells on the dorsal side of the embryo and elongated cells on the ventral side of the embryo. Insets show the dorsal and ventral side of the forming blastopore lip with higher magnification. See **Supplemental Figure 5.A** for schematic. **B.** Dorsal-lateral optical plane view shows distinct anillin staining patterns in different parts of the forming dorsal blastopore lip. Consistent with the optical cross-section view, anillin has a membrane proximal localization in elongated and both elongated and apically constricted cells at the dorsal blastopore lip. In the younger and more vegetal region of the lip, the localization of anillin is punctate and in the older more animal lip the localization is contiguous and smooth.

**Figure 8.**
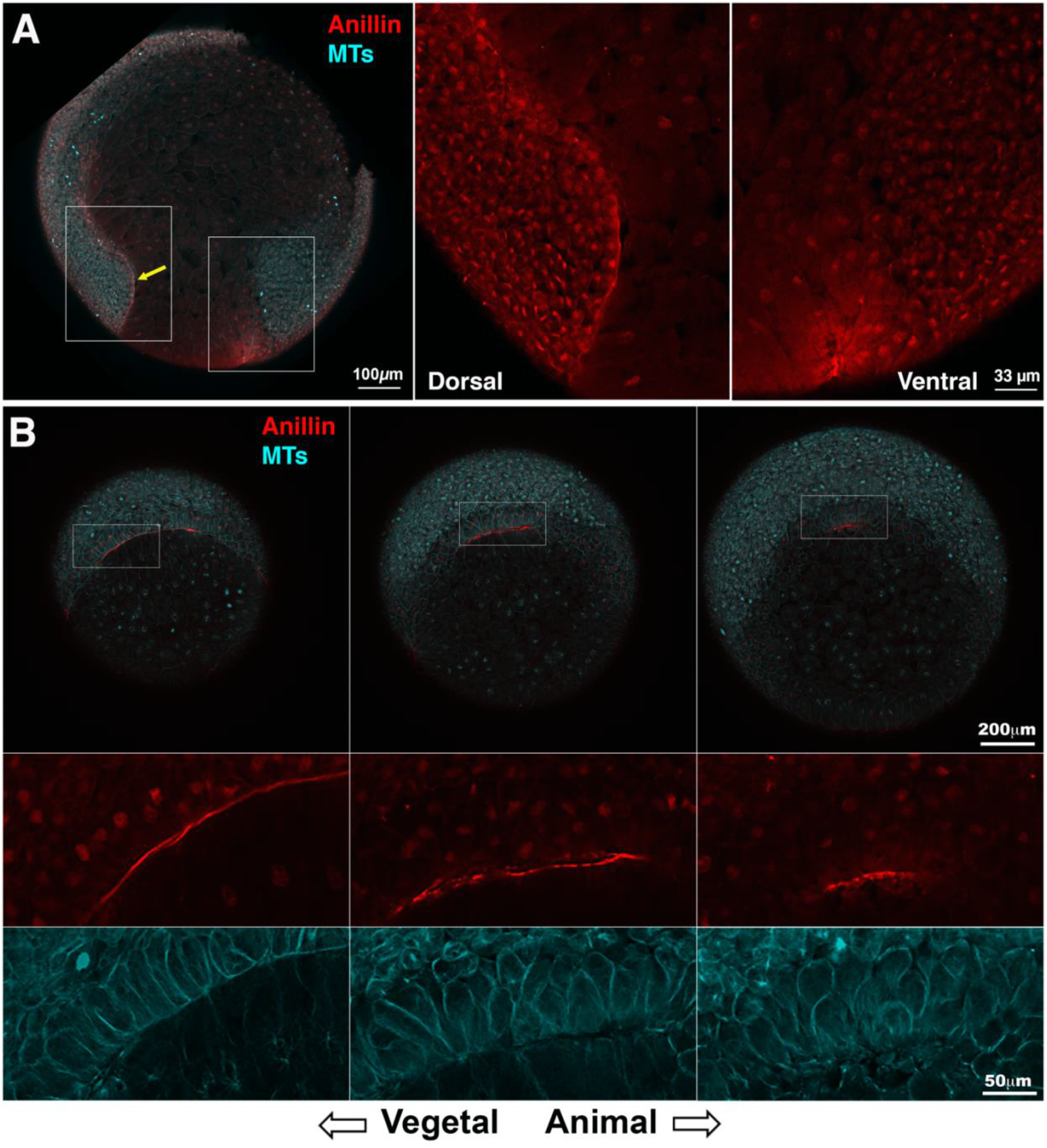
Whole embryo fixed immunofluorescence against tubulin and anillin after dorsal lip maturation enables visualization of the forming archenteron. **A.** Animal-vegetal optical plane view of a Stage 11 embryo shows membrane proximal anillin staining along the mature dorsal and immature ventral blastopore lips. Insets show high magnification. The anillin staining is along the animally extending archenteron (yellow arrow). See **Supplemental Figure 5.B** for corresponding schematic. **B.** Dorsal-lateral optical plane views show the differences in anillin staining and cell shape in different regions of the mature lip and archenteron. More vegetally the anillin staining is often in regions of packed columnar or square cells. In these regions the anillin appears as a line basally. As the planes move animally, the anillin staining is more spotted, and the cells are various shapes that are neither square shaped nor spherical. When the whole embryo views of anillin staining are compared, the span of anillin staining get shorter as the cross sections progress from more vegetal to more animal. This reveals that the archenteron is formed and is moving animally.

The cells on the dorsal side of the archenteron contact involuted chordal mesoderm that is expected to be expressing Wnt11 family ligands. Another possibility is that there is no Wnt11 family mediated signaling in the tissue itself, but that both maturation of bottle cells into a contiguous lip and extension of the archenteron require Wnt11 family dependent forces acting on the deep tissues above the blastopore. Interestingly, non-canonical Wnt signaling proteins including Shroom2 [38], Vangl2 [50] and Strabismus [51] have all been reported to play roles in epithelial cells undergoing shape changes related to gastrulation indicating that PCP signaling is occurring in the epithelium itself. If this were the case, there should be a tight relationship between PCP activated Rho-signaling and anillin. Further molecular perturbation experiments would be required to test the function of anillin in this process and in gastrulation morphogenesis more generally. In some cases, such experiments would be complicated by the fact that anillin is essential for cell division. Much has been made of the fact that cell division appears to have stopped in the chordal mesoderm by the middle of gastrulation; however, cell divisions do occur elsewhere in the embryo [52, 53]. Even in the absence of an understanding of anillin’s mechanistic role in gastrulation, anillin localization should be a useful marker for future research on the morphogenetic pathways responsible for blastopore lip maturation and archenteron formation.

## Acknowledgements

The authors acknowledge Leonid Peshkin and Jenny Gallop for the gift of the embryo holders used for the inverted external imaging, Rachael Jonas-Closs for exceptional Xenopus husbandry, and the Nikon Imaging Center at Harvard Medical School for microscopy and image analysis resources. Funding to TJM from GM131753 and to MWK from HD073104.

## Author Contributions

*Elizabeth Van Itallie:* Conceptualization, Investigation, Writing – Original Draft, Writing – Review & Editing; *Christine M. Field:* Conceptualization, Investigation, Writing – Review & Editing; *Timothy J. Mitchison:* Resources, Writing – Review & Editing, Funding acquisition. *Marc W. Kirschner:* Writing – Review & Editing, Funding acquisition.

## Methods

### Normal Table Illustrations

The external view vegetal pole Normal Table drawings in Supplemental Figure 5 are from Xenbase (www.xenbase.org; RRID:SCR_003280) [54]. Illustrations © 2021 Natalya Zahn, CC BY-NC 4.0 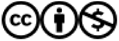

### Frog Use

All handling of Xenopus laevis adults was done in accordance with HMS IACUC protocol IS00001365. Females were injected with 500 units of human chorionic gonadotropin (HCG) to stimulate oocyte maturation and laying. All embryos result from in vitro fertilization (IVF). Embryos were cultured in 0.1x MMR at 18C unless otherwise noted [55].

### Morpholinos

All morpholino injections were done at the 2-cell stage, and for all experiments morpholino was injected into both blastomeres. The morpholino amount noted for each experiment is the total amount of morpholino injected into the embryo. During injections, embryos were cultured in 0.5x MMR, 4% ficoll solution (Ficoll PM 400, Sigma). Injected embryos and fertilization matched sibling control embryos were cultured in 0.5x MMR, 4% Ficoll until they reached Stage 9 when they were washed into 0.1x MMR. The control morpholino is the Standard Control Morpholino sold by GeneTools. The sequence of the Wnt11b morpholino (5’ to 3’) is CCAGTGACGGGTCGGAGCCATTGGT. The sequences of the Wnt11 morpholinos (1) and (2) are CTGTATCCAAAGAGAGTTCCGAGGT and ATGAAAAGATCCTGGATTGGCAGTC, respectively. Morpholinos were resuspended in water, aliquoted, and stored at -80°C. They were heated at 60°C for 10 minutes before combining at the appropriate concentrations for injection [56]. For the Standard Control morpholino, 1 picomole equals 8.3 ng. For all of the Wnt11 family morpholinos, 1 picomole equals approximately 8.5 ng.

### Still image phenotype quantification

For all quantification of end of gastrulation and tailbud stage phenotypes, the staging is done using the un-injected sibling embryos. For the blastopore closure analysis from the time-lapse imaging, the frame corresponding to Stage 10.5 for all of the sibling embryos was determined and that same frame was used as the starting point for the injected embryos.

### Movies

Custom made plastic inserts were coated with poly-hema solution and placed in the chambers of Lab-Tek II eight chambered #1.5 german cover glass systems (#155409). Chambers were filled with 0.75 mL of 0.1x MMR, and three embryos were place in each well. Their vegetal poles naturally oriented towards the objective lens due to gravity. The room temperature was 19°C. Time lapse images of gastrulation were collected with a Nikon Eclipse Ti2 Inverted microscope using a 4x objective lens, Zyla sCMOS camera, and Nikon Elements software. The embryos were illuminated obliquely from below using a diode light source mounted on a flexible arm. This resulted in darkfield illumination with contrast provided by the cortical pigment granules and the color differences between cell surfaces and cell-cell interfaces. The relatively weak dark-field diode illumination did not significantly warm the embryos or perturb the embryos, so a shutter was not required. Eleven Z-plane images were acquired for each embryo every five minutes with an acquisition time between 500-700 ms. The Z-stacks were processed with Nikon Elements Extended Depth of Field (EDF). Kymographs were made with the FIJI software kymograph program (https://fiji.sc/).

### Immunofluorescence

Embryos were fixed and stained as previously described [57]. Briefly, embryos were fixed in 50 mM EGTA, ∼ pH 6.8, 10% H2O, 90% methanol for at least 24 hours at room temperature with gentle shaking. They were then be stored at 4°C. Prior to staining, embryos were rehydrated in a series of steps: 25%, 50%, 75% and 100% TBS (50 mM Tris, pH 7.5, 150 mM NaCl/methanol) for 30-45 min per step, with gentle shaking. Embryos were bleached overnight in a solution of 1% H2O2, 5% formamide, 0.5x SSC (75 mM NaCl and 8 mM sodium citrate, pH 7). Embryos were rinsed 3x in TBS and then incubated with directly labeled antibodies for at least 24 hours at 4°C with very gentle rotation. Antibodies were diluted in TBSN (10 mM Tris-Cl, pH 7.4, 155 mM NaCl, 1% IGEPAL CA-630), 1% BSA, 2% fetal calf serum (FCS) and 0.1% Sodium Azide. After antibody incubation, embryos were washed in TBSN for approximately 48 hours (with several solution changes), and then washed 2X in TBS. Prior to clearing (see below), embryos were dehydrated in four steps depending on the clearing agent used. For Murray Clear (66% benzyl benzoate, 34% benzyl alcohol), the steps were 30%, 50%, 75%, and finally 100% methanol in TBS. For Ethyl cinnamate, the steps were 30%, 50% and 75%, and finally 100% 1-propanol in PBS [58]. Each dehydration step is approximately 45 minutes. After the embryos were transferred to a clearing solution, they were mounted in clearing solution filled circulator holes in metal slides (∼1.2 mm thick) that were capped by coverslips on both sides. Murray Clear and ethyl cinnamate performed similarly with respect to clearing and imaging, but the latter may present less toxicity risk.

### Microscopy Immunofluorescence

Confocal Z-stack images were obtained at the Nikon Imaging Center at Harvard Medical School. Fixed embryos were imaged using a Nikon Ti-E inverted microscope with a Nikon A1R point scanning confocal head using 10x and 20x dry objectives and NIC Elements acquisition software.

### Antibodies

Antibodies raised against *Xenopus laevis* anillin [32] and alpha-tubulin (Sigma #T6074) were labeled on a column with Alexa-568, or Alexa-647 dyes (Life Technologies, NY) as previously described [57].

---

**Supplemental Figure 1.**
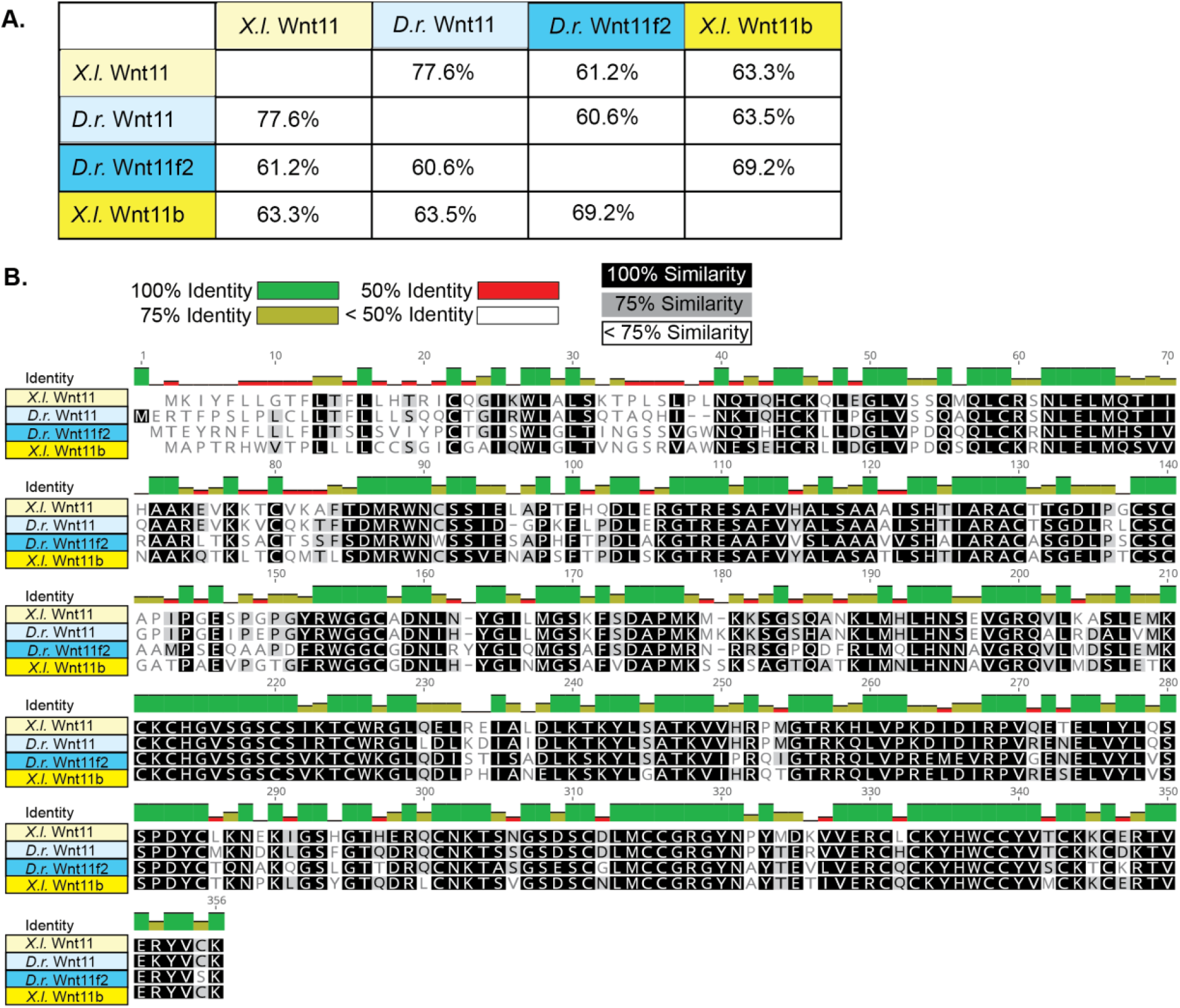
The *Xenopus laevis* and *Danio rerio* (zebrafish) genomes both encode two Wnt11 family proteins. **A.** *Xenopus laevis* Wnt11 and Wnt11b are more similar in protein sequence identity to their *Danio rerio* homologs (77.6%, 69.2%) than they are to each other (63.3%). **B.** Multiple sequence alignment of the *Xenopus laevis* and *Danio rerio* Wnt11 family members shows that despite the closer homology between the Wnt11s and Wnt11b/Wnt11f2, there are regions of 100% similarity between all four proteins across the entire protein sequence. The first ∼20 amino acids are a signal peptide that is cleaved in the forming of mature proteins. (Geneious Prime 2020; https://www.geneious.com)

**Supplemental Figure 2.**
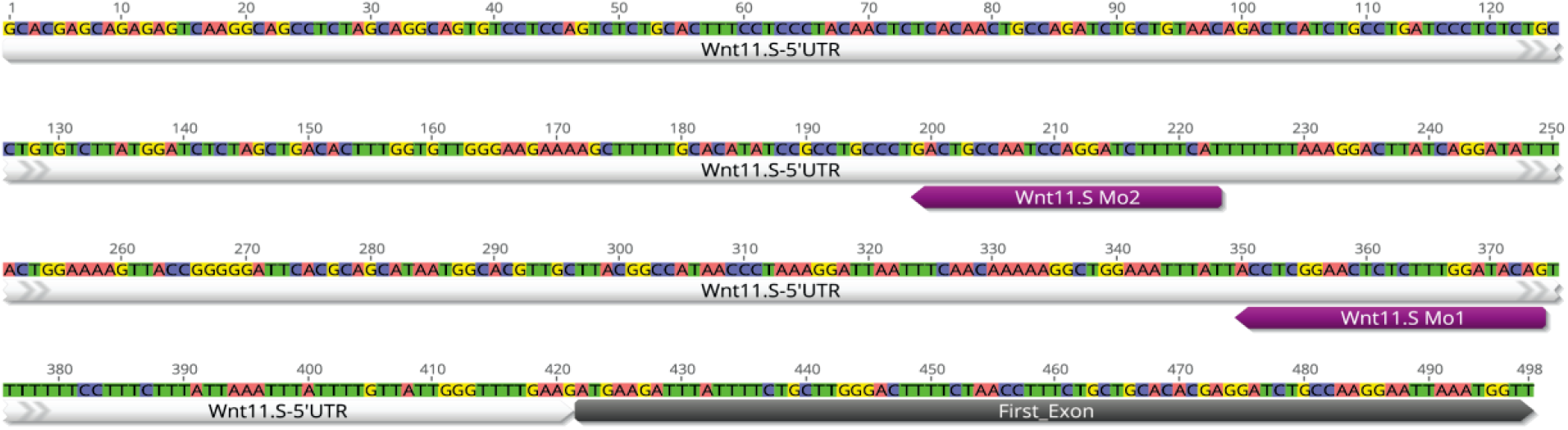
The location of the targets of Wnt 11 Mo1 and Wnt11 Mo2 in the 5’UTR of Wnt11.S. (Geneious Prime 2020; https://www.geneious.com)

**Supplemental Figure 3.**
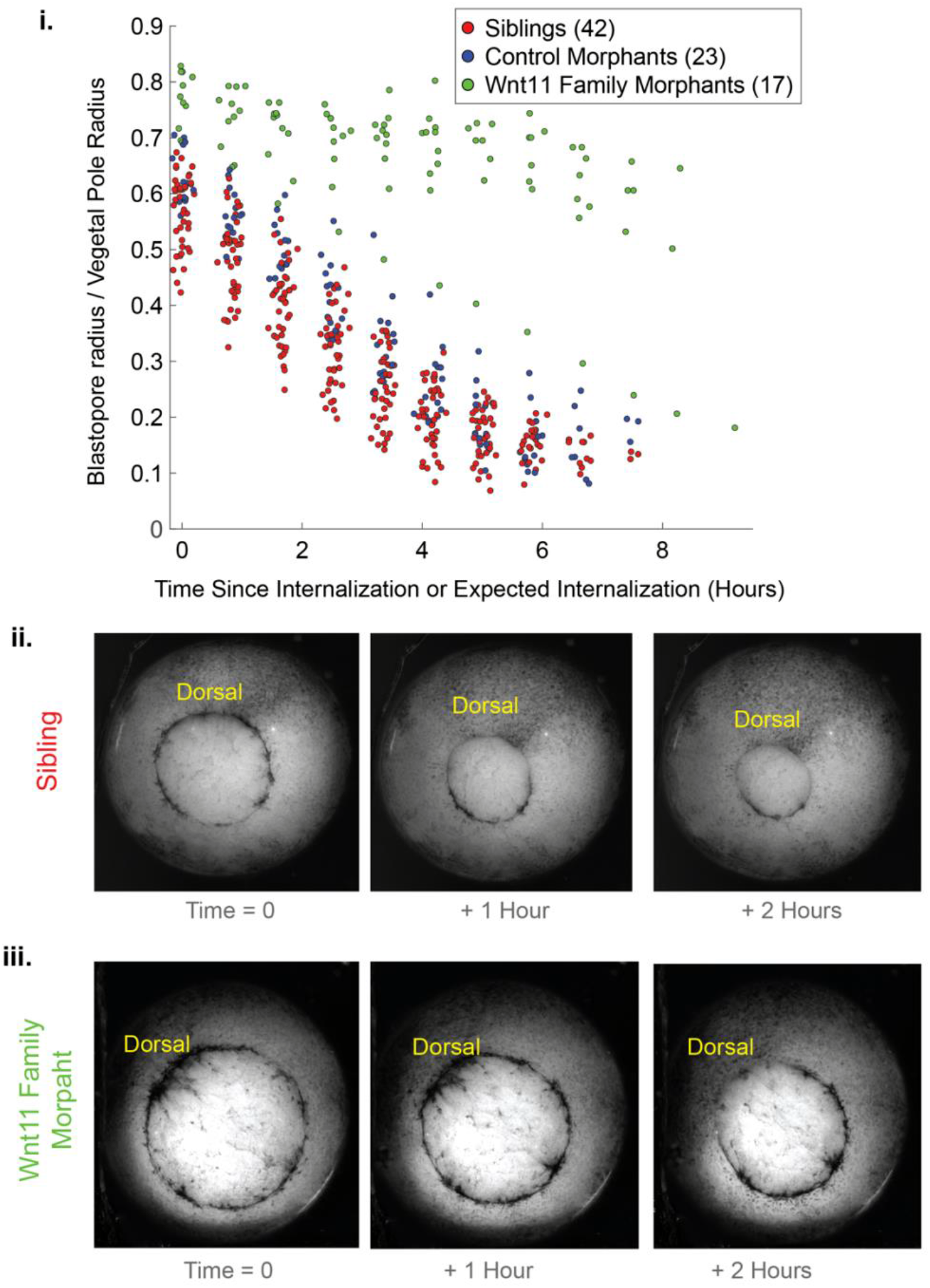
Wnt11 family morphant embryos have larger blastopores at the time when the dorsal lip matures in sibling morphants; and Wnt11 family morphant embryos exhibit less blastopore closure during the second half of gastrulation than sibling and control morphant embryos. **(i)** Quantification of the decrease in normalized blastopore radius for the sibling, control morphant, and Wnt11 family morphants during gastrulation. Quantification starts when sibling embryos form a mature blastopore lip (as determine by inspection of movie frames). For control morphant and Wnt11 family morphant embryos, quantification begins at the expected time of lip maturation as determined by the mean of the matched sibling maturation times. Wnt11 family morphants have larger blastopore radii when quantification begins and show only limited blastopore closure afterwards. In addition to Wnt11 family morphant phenotypes, control morphant embryos have on average larger blastopores at the expected time of lip maturation than siblings. This is consistent with the expected delay of gastrulation caused by any morpholino injection. However, the blastopores of control morphants do close over a similar time period as the sibling embryos. **(ii, iii)** Still images for a sibling embryo and Wnt11 family morphant embryo at the beginning of quantification, and one and two hours later. Progressive closure of the blastopore is shown in the sibling embryo. For the Wnt11 family embryo the blastopore has formed by the expected lip maturation time, but the dorsal bottle cells are entirely visible. At the one hour timepoint the blastopore is slightly smaller, but the dorsal lip has still not matured. After two hours the dorsal vegetal material has extruded above the dorsal lip instead of being internalized.

**Supplemental Figure 4.**
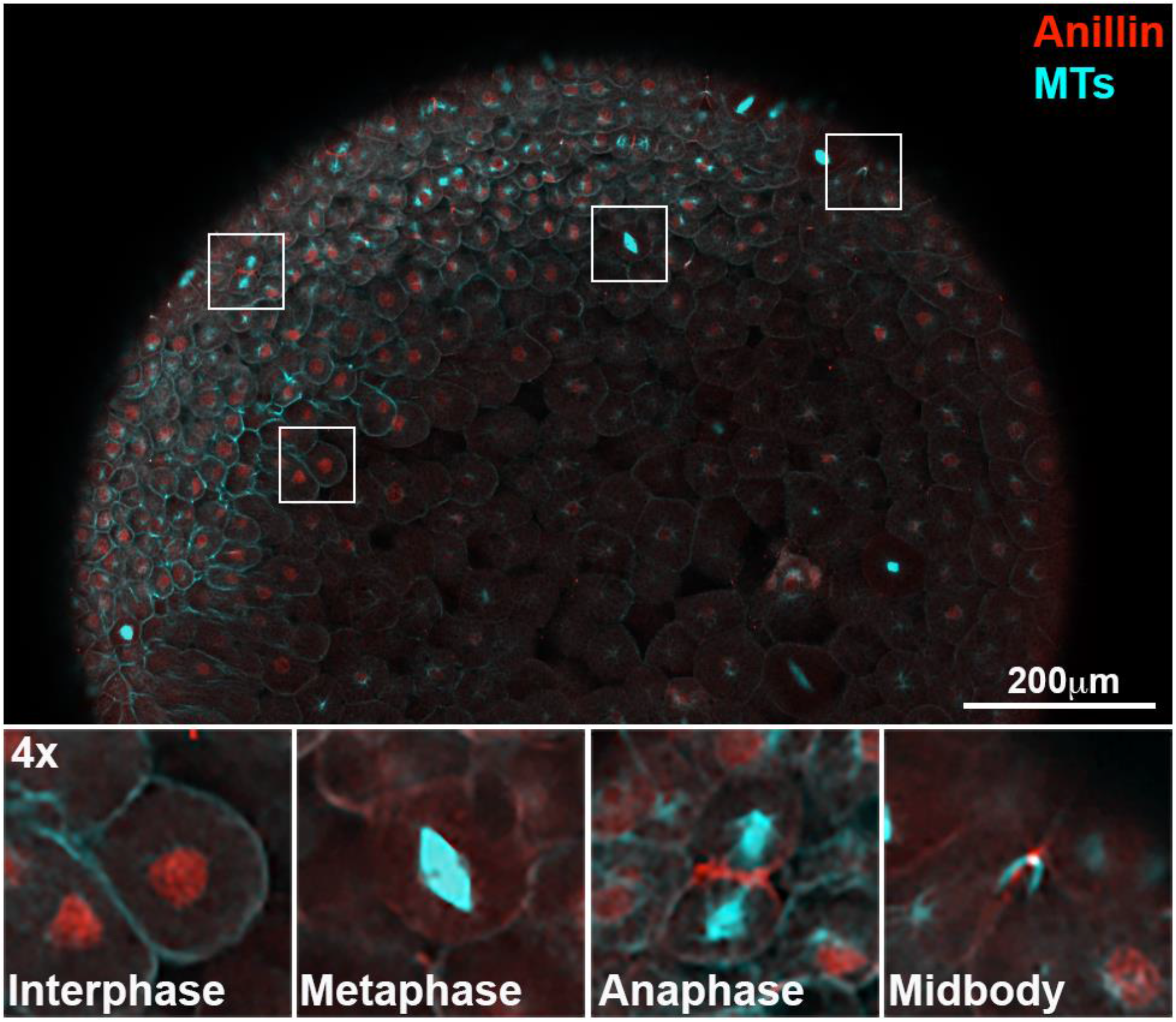
Antibodies against anillin (red) and microtubules (cyan) have the expected localizations related to the cell cycle. Inset panels show examples of cells in interphase (anillin in the nucleus), metaphase (microtubule spindle), anaphase (microtubule spindles separating on either side of the anillin localized cleavage furrow, and the end of cytokinesis (anillin in the microtubule midbody structure).

**Supplemental Figure 5.**
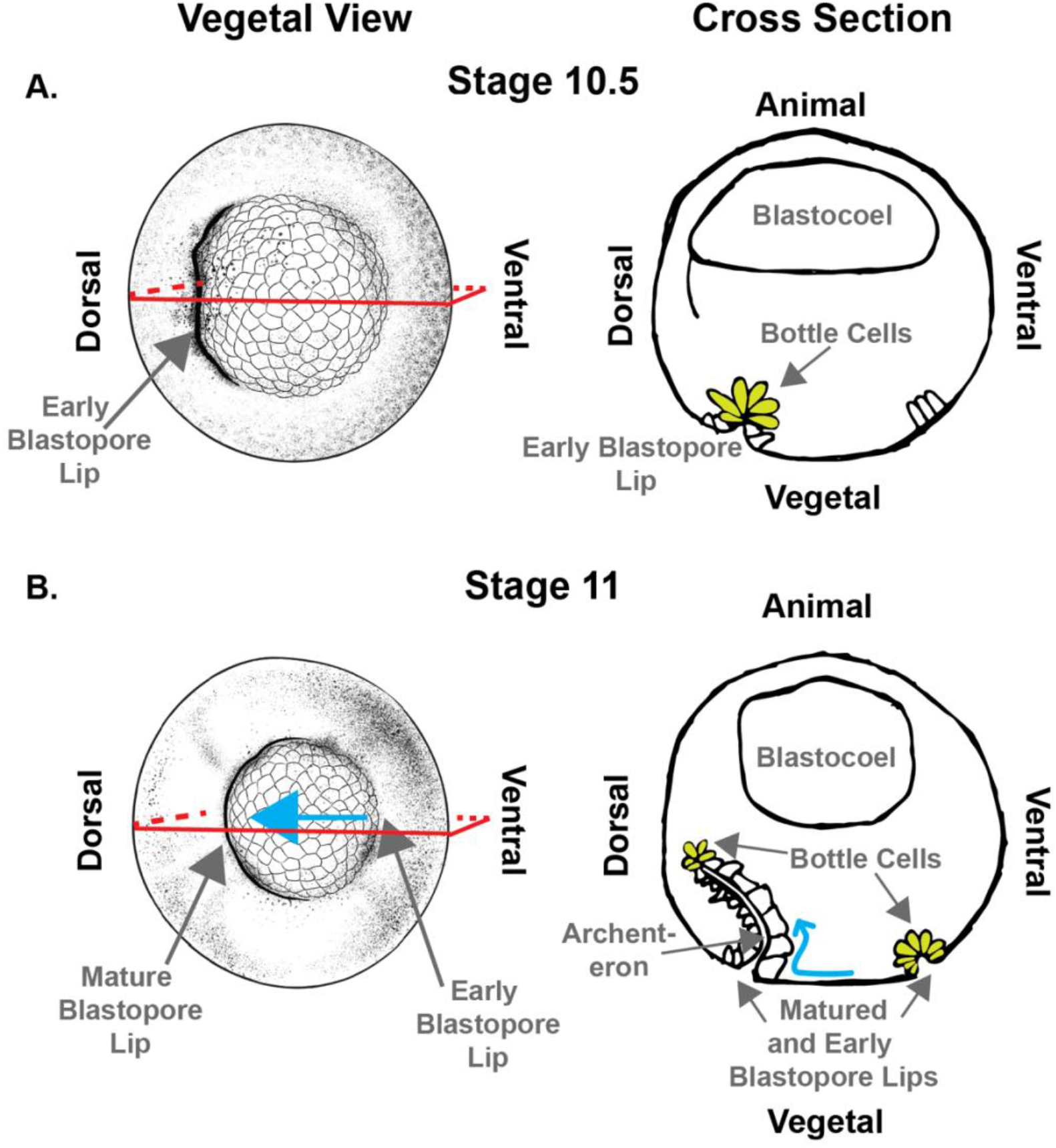
Drawings and schematics showing two views of embryos at Stages 10.5 and 11. On the left, external-vegetal views, and on the right internal cross-section views. The schematics show the locations of the structures discussed in the main text: bottle cells, blastopore lips, and the archenteron. The red box shows the relationship of the vegetal view drawing to the cross-section schematic before a 90-degree upwards rotation. **A.** At Stage 10.5 the early blastopore lip has formed on the dorsal side of the embryo. The lip is dark in color because bottle cell apices are at the lip. There are elongated pre-bottle cells where the dorsal lip will form. **B.** At Stage 11 the dorsal blastopore lip is mature. The external color of the mature dorsal blastopore lip has changed because the bottle cells have left the blastopore lip and moved animally. The bottle cells are at the animal extent of the archenteron which is lined by the epithelial cells that were formally above and below the early blastopore lip. Bottle cells have formed on the ventral side of the embryo resulting in an early blastopore lip. Vegetal rotation (dorsal animal movement of the vegetal endoderm; blue arrows) is occurring internally as early as Stage 10, but it is clearly visible from movies of the exterior of the embryos at Stage 11. Cross-section drawings are based on based on drawings by Peter Hausen and Metta Riebesell [1].

